# Kernel Matrix Completion with Topological and Spectral Features for Multi-Modal Classification

**DOI:** 10.64898/2026.04.19.713528

**Authors:** Ela Mae Riñon, Maria Vivien Visaya, Rachelle Sambayan

## Abstract

Kernel methods offer a robust framework for integrating multi-modal datasets into a unified representation, thereby facilitating more comprehensive data interpretation. In the presence of incomplete datasets, multiple kernel learning is employed to enhance the efficiency of data completion and integration. We investigate kernel-based approaches to address the incomplete-data problem with applications to yeast protein data. Biological data such as yeast proteins can be represented through multiple modalities, including gene expression profiles, amino acid sequences, three-dimensional structures, and protein interaction networks. We introduce a computational pipeline based on kernel matrix completion, in which topological data analysis (TDA) and persistent spectral analysis are incorporated into the classification setting. TDA captures geometric structure across scales while spectral descriptors reflect connectivity patterns through Laplacian eigenvalues. Kernel, topological, and spectral descriptors are used with support vector machines to discriminate between membrane and non-membrane yeast proteins. Empirical results show that the combined pipeline improves both kernel completion accuracy and ROC performance relative to baseline kernel-only approaches. The best-performing configuration achieves an ROC score of 0.8632 using the average of three kernels augmented with TDA features. These results demonstrate competitive performance relative to strong kernel-based baselines under incomplete data conditions. The proposed approach provides a unified approach for learning from incomplete heterogeneous data while enriching kernel representations with geometric and spectral information.

## 1. INTRODUCTION

Protein data are naturally represented through multiple modalities, including sequence, expression, structural, and interaction-based information. Recent work in pattern recognition has increasingly focused on multi-modal and multi-view data, where complementary information across sources must be integrated [17, 19]. Learning from such data is further complicated by missing observations, which disrupt the similarity structure used in downstream analysis. We consider the problem of learning from incomplete heterogeneous data using similarity-based representations. From a pattern recognition perspective, this corresponds to learning structured representations from incomplete data, where multiple sources induce complementary similarity relationships.

Protein classification methods aim to categorize proteins by examining their global sequence and structural similarities. However, proteins, while fundamental to nearly all physiological processes, rarely function in isolation. A single protein often participates in multiple cellular roles across various biological processes, from the molecular to the systemic level. This multifunctionality introduces challenges in modelling protein relationships across different biological contexts. This motivates the development of classification methods that capture the underlying biological relationships among proteins.

Work in protein representation learning highlights the need for methods that incorporate structural, functional, and interaction-based information [20, 33]. In bioinformatics, recent studies have increasingly focused on protein-protein interactions, developing methods capable of analyzing large-scale genomic and proteomic data. The work in [34] studies protein interactions using network-based representations, while [8] uses sequence-derived features for protein analysis, and [1] focuses on large-scale genomic and proteomic data integration. Recent large-scale learning approaches have also been used to model protein data across diverse sources [26, 30]. These challenges are closely related to multimodal data fusion, where heterogeneous sources must be combined in a consistent representation [6]. These approaches extract relationships that go beyond sequence and structural similarity by incorporating multiple protein modalities such as gene expression profiles, interaction networks, and sequence data [1]. As biological datasets grow in size and complexity, the heterogeneity of protein properties and functions further complicates the development of computational methods that can effectively capture these interactions.

Kernel methods offer a natural way to analyze such data by mapping them into higher-dimensional feature spaces, where underlying patterns can be more easily uncovered [2]. A common strategy is to define protein kernels that quantify similarity based on specific characteristics. Ben-Hur *et al*. [3] extended this approach by introducing pairwise kernels, which evaluate similarity between protein pairs rather than individual proteins, improving predictive performance when combined with support vector machines. Despite their effectiveness, kernel methods face limitations, including sensitivity to incomplete data and the computational cost associated with high-dimensional representations [4, 10]. In related settings, incomplete multi-view learning has been studied to handle missing observations while preserving structural relationships across data sources [11].

Graph-based representations are central to modelling biological systems, particularly protein–protein interaction networks. Recent work in graph representation learning has shown strong performance in modelling protein interaction data [16, 28]. However, recent graph and deep learning approaches achieve strong performance but typically assume complete data [15]. Learning from incomplete kernel matrices has been studied in the context of kernel completion, where the goal is to reconstruct missing similarities while preserving the underlying structure of the data [29]. However, these approaches typically do not consider how kernel completion influences downstream structural feature extraction. The focus in this work is on understanding representation structure under incomplete similarity information, rather than optimising end-to-end predictive performance.

Recent work in pattern recognition has increasingly incorporated geometric and topological representations, emphasizing the importance of structured representations for complex data [12, 22] Topological Data Analysis (TDA), particularly persistent homology, offers a complementary perspective by focusing on the shape of data rather than individual observations. It identifies features such as clusters and loops that persist across multiple scales [5], and summarises them through compact representations such as persistence diagrams and landscapes.

Topological data analysis has been increasingly applied in biological and biomedical settings to capture structural patterns beyond pairwise similarities. Recent surveys highlight its effectiveness in analyzing complex molecular and omics data [31]. In machine learning, persistence-based representations such as persistence diagrams and their vectorisations have been incorporated into learning models to extract discriminative topological features [13, 21]. Spectral approaches based on Laplacian operators have also been extended to topological settings, where persistent Laplacians give a multi-scale description of structural properties. These methods have been applied to biomolecular data and capture information not reflected in standard topological summaries [24].

Several recent studies have demonstrated the value of combining topological and spectral information with machine-learning methods for analyzing biological data. More broadly, recent work has explored the use of topological features in graph and representation learning pipelines, further highlighting the role of multi-scale structure in pattern recognition tasks [35]. For instance, [32] combines persistent spectral embeddings with sequence and structure representations to model protein fitness landscapes. These developments support the integration of spectral and topological descriptors within a unified representation. Empirical studies suggest that persistent spectral features can provide complementary or improved predictive performance relative to persistent homology in biomolecular learning tasks [23]. Related approaches combining spectral descriptors with machine learning have also been used in protein–ligand binding prediction and protein solubility analysis [36]. These observations motivate combining kernel, spectral, and topological features within a single representation. From a pattern recognition perspective, this corresponds to learning from incomplete structured data, where each modality induces a similarity-based representation. While recent advances in deep and graph-based learning have shown strong performance in biological applications, these approaches typically assume complete data and do not explicitly address the reconstruction of missing similarities or the incorporation of multi-scale geometric descriptors. In contrast, the approach proposed in this work integrates kernel-based representations with topological and spectral descriptors to construct features under incomplete data conditions.

Functional complexity in proteins cannot be captured by sequence or structure alone, particularly when data are incomplete or originate from multiple sources. We address this by constructing a classification approach that brings together complementary topological, spectral, and similarity-based information, and examine how these representations jointly reveal structure not captured by individual modalities.

This study provides a systematic assessment of how kernel learning, persistent homology, and persistent spectral analysis enhance the classification of yeast membrane and non-membrane proteins, and how each component contributes to the representation and influences predictive performance [25]. The main contribution of this work is a unified approach for learning from incomplete heterogeneous data, where kernel matrix completion is combined with topological and spectral feature extraction to produce representations that capture both similarity structure and multi-scale geometry. Rather than treating completion as a preprocessing step, the proposed approach incorporates it directly into the feature construction pipeline, allowing downstream descriptors to reflect the reconstructed structure of the data. We demonstrate the effectiveness of this approach on a protein classification task, where the integration of kernel, topological, and spectral information improves predictive performance over standard kernel-based baselines.

The contribution of this work is not in introducing a new topological or spectral descriptor in isolation, but in how these components are integrated within the learning pipeline. In particular, kernel matrix completion is treated as part of the representation construction, rather than as a preprocessing step, so that the reconstructed similarity structure directly influences downstream topological and spectral features. This interaction between completion and multi-scale feature extraction is not explicitly considered in existing kernel completion or topological learning frameworks. The proposed approach therefore provides a structured way to analyze incomplete data where similarity, topology, and connectivity are coupled within a single representation.

## 2. METHODOLOGY

The proposed methodology differs from standard kernel-based pipelines in that feature extraction is performed after reconstructing incomplete kernel matrices, rather than on fully observed data. This allows the topological and spectral descriptors to reflect the recovered similarity structure, which is particularly relevant in settings where data from multiple sources are incomplete. The overall pipeline therefore combines kernel completion, topological data analysis, and persistent spectral analysis within a single representation.

To assemble the information carried by different representations of yeast proteins, we implement a workflow that brings together kernel completion, topological descriptors, and persistent spectral features. We begin by reconstructing incomplete kernel matrices using kernel matrix completion, providing a stable similarity structure on which subsequent analyses are built. Topological data analysis is then applied to quantify the multiscale topology encoded in the completed matrices, while persistent spectral analysis captures harmonic information derived from the combinatorial Laplacian. Finally, we combine the resulting kernel-based, topological, and spectral features within an SVM classifier to distinguish membrane and non-membrane proteins. This section describes the data, the feature extraction procedures, and the experimental setting used in the study. Figure 1 illustrates the feature extraction process through kernel matrix completion, topological data analysis (TDA), and persistent spectral analysis (PSA). The input protein data consist of multiple heterogeneous sources, where each protein *P*_*i*_ is represented by features derived from sequence information, gene expression measurements, and protein-protein interaction (PPI) data. The different marker shapes (circles, stars, triangles) illustrate distinct types of raw features originating from these data modalities. The following stages are then performed: (1) These features are used to construct partially observed kernel matrices, which are completed using the PCAPSA-MKMC method to obtain a full similarity matrix. (2) The completed kernel matrix is then used to compute topological features (via persistent homology) and spectral features (via persistent spectral analysis). (3) Kernel-derived features, topological features, and spectral features are combined to form a unified representation of each protein. (4) A machine-learning classifier (SVM) is trained on the feature set to distinguish membrane proteins from non-membrane proteins.

**Figure 1.**
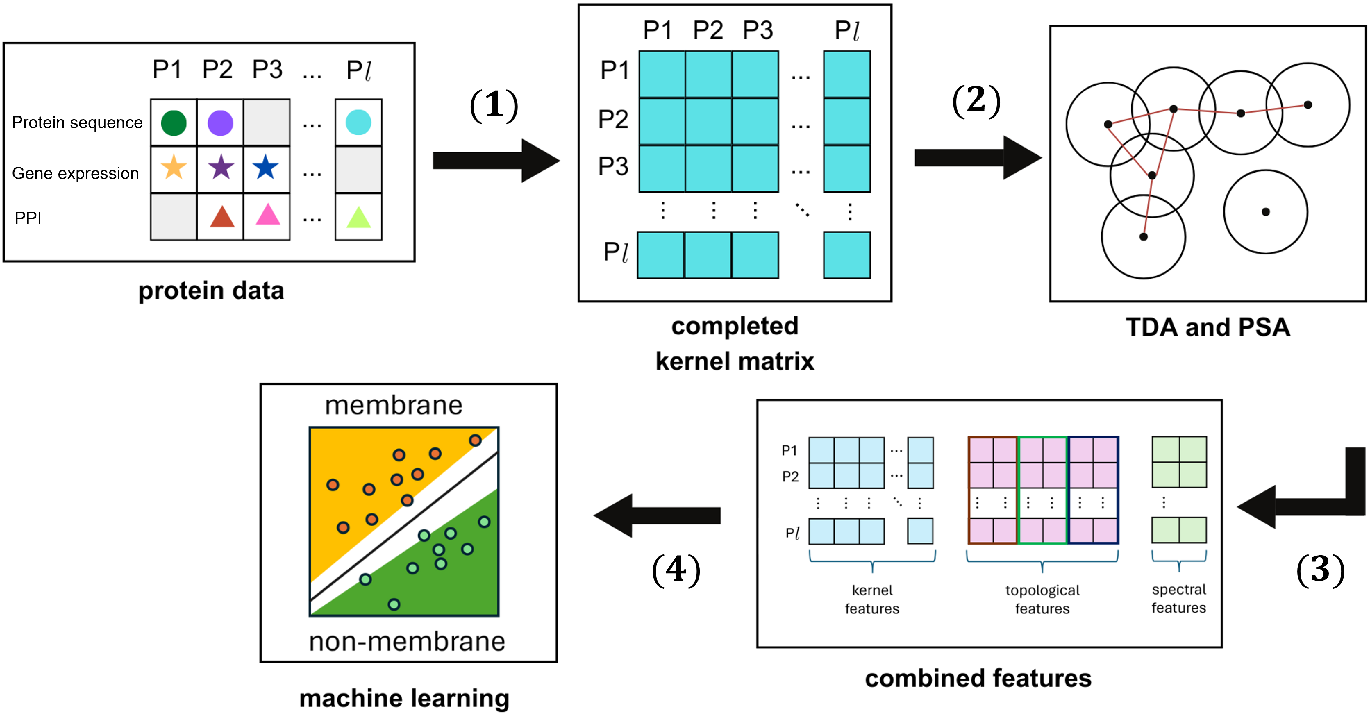
Overview of the proposed framework. Protein data from multiple sources are used to construct partially observed kernel matrices that are completed using PCAPSA-MKMC. Topological (persistent homology) and spectral (persistent spectral analysis) descriptors are then extracted and combined with kernel features for SVM-based membrane protein classification.

Following the overall workflow shown in Figure 1, we describe each of the three main methods in detail.

### Kernel matrix completion

We represent the *l* proteins using an *l* × *l* kernel matrix, *Q*, which is a positive definite matrix where the (*i, j*)−th entry denotes the similarity between the *i*th and *j*th proteins. In cases where data for some proteins are unavailable, the corresponding entries in the kernel matrix remain missing. These missing entries prevent the application of machine learning algorithms. To address this issue, we employ the Principal Component Analysis-based Mutual Kernel Matrix Completion (PCA-MKMC) algorithm [15] developed by Rivero and Kato. This parametric model for mutual kernel matrix completion facilitates the completion of the matrix. The resulting completed kernel matrix yields unique descriptors for each protein, referred in this study as **kernel features**.

### Topological Data Analysis

After obtaining the completed kernel matrix, the next step is to convert it into a distance matrix for compatibility with TDA applications. We perform filtration by creating balls around each data point with an expanding radius. From this filtration, we construct a simplicial complex— a structure composed of simplices such as points, edges, and triangles—that captures the topological features of the data. Once the simplicial complex is built, we proceed with persistent homology, monitoring the birth and death of topological features. The results are visualized through a persistence diagram, where each birth–death pair obtained from the filtration represents a feature and plotted on the *x*− and *y*−axes, respectively. The persistence diagram is transformed into a persistence landscape {*λ*_*m*_(*t*)}_*m*≥1_, where *t* is the filtration parameter and where each *λ*_*m*_ : ℝ → ℝ is the *m*-th largest value of the collection of piecewise-linear functions induced by the birth–death intervals. From the resulting landscape functions, we compute their *L*^*p*^-norms

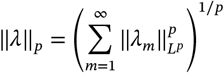

which yield a finite-dimensional summary of the persistence information. These norms are used as an additional identifier for each protein, and we refer to them as the topological features of the protein.

### Persistent Spectral Analysis

The filtration process systematically transforms the point cloud into a sequence of chain complexes, generating a family of simplicial complexes and their associated chains. From these, we construct persistent combinatorial Laplacian matrices, which encode information about connectivity and the underlying structure of the space. Spectral features are derived by computing the eigenvalues and eigenvectors of these Laplacians at each filtration scale. In particular, tracking the evolution of zero eigenvalues captures persistent harmonic modes, which encode robust topological invariants [25]. Extracting these spectra over the filtration constitutes PSA, providing complementary structural information to persistent homology.

## 3. EXPERIMENTAL SETTINGS

This study develops a framework for classifying yeast proteins as membrane or non-membrane. This section begins with the presentation of protein data and data sources, followed by feature extraction using kernel methods, topological data analysis, and persistent spectral analysis. Each protein is characterized by a distinct set of extracted features, which are then combined and used as inputs for machine learning algorithms. Specifically, we use support vector machines (SVMs) for classification.

### 3.1. Protein Data and Sources

We consider *Saccharomyces cerevisiae* (brewer’s or budding yeast), which encodes 6,112 proteins. Subcellular location annotations from the MIPS Comprehensive Yeast Genome Database (CYGD) cover 2,318 proteins, including 497 membrane-associated and 1,821 non-membrane proteins; the remaining 3,794 are unlabeled [1]. For each protein, we consider three data sources that capture distinct information relevant to the classification task: **protein sequences**, which capture structural and physicochemical properties; **gene expression profiles**, which reflect functional activity across experimental conditions; and **protein–protein interaction (PPI) networks**, which model connectivity patterns within the cellular system.

#### Protein sequences

Sequence similarity to known membrane proteins is often indicative of shared localization [7]. In addition, structural motifs such as alpha-helices and beta-sheets, commonly observed in membrane proteins due to their interaction with the lipid bilayer, yield useful discriminative features [14].

#### Gene expression

Expression profiles capture compartment-specific activity. Membrane proteins are typically associated with the plasma membrane, endoplasmic reticulum (ER), or mitochondria [18], while non-membrane proteins show patterns linked to intracellular regions such as the cytosol or nucleus, reflecting their diverse functional roles. These contrasting patterns make gene expression highly informative for classification.

#### Protein–protein interactions

PPI networks further differentiate membrane proteins, as hydrophobic regions promote preferential interactions [14]. Transmembrane proteins are also frequently involved in signalling pathways, reflecting their roles in cellular communication and transport [18]. These interaction patterns highlight functional roles distinctive to membrane proteins.

### 3.2. Protein Feature Extraction

To represent each protein in a manner suitable for classification, we extracted features from the three data sources described in Section 3.1 and shown in Table 1. Each data source contributes information, and features were derived using three distinct approaches: kernel-based similarity measures, topological data analysis, and persistent spectral analysis.

**Table 1.** Kernel matrices constructed from different data sources and the kernel functions used for their construction.

| Kernel matrix | Data source | Kernel function |
| --- | --- | --- |
| $Q^{(1)}$ | Protein sequences | Linear kernel |
| $Q^{(2)}$ | Gene expression | RBF kernel |
| $Q^{(3)}$ | Protein interactions | Diffusion kernel |
| $Q^{(c)}$ | Combined data sources (1) | |

#### 3.2.1. Kernel-Based Features

Kernel-based features capture pairwise similarities among proteins by applying kernel functions to each data type: a linear kernel for sequence data, a radial basis function (RBF) kernel for gene expression, and a diffusion kernel for PPI networks. Choosing an appropriate kernel function often requires understanding the underlying data structure and the problem domain. These choices align with prior studies [1] which have identified them as well-suited for protein analysis. Computing the similarity between two proteins based on a specific data source and its corresponding kernel function results in a symmetric, positive semi-definite matrix.

##### *Q*^(1)^: Linear kernel matrix derived from protein sequences data

The similarity between two distinct protein sequences is measured using the **BLAST (Basic Local Alignment Search Tool)**, which evaluates sequence alignment based on a scoring matrix. Specifically, BLAST uses the **BLOSUM62 substitution matrix** to assign similarity scores to aligned residues. For a given pair of proteins *i* and *j*, the final similarity score is computed as the sum of individual substitution scores, adjusted by penalties for gaps introduced during alignment. Since BLAST score matrices are not guaranteed to be positive semi-definite, each protein is instead represented as a vector of similarity scores relative to all other proteins. A valid kernel matrix *Q*^(1)^ is then constructed by computing the Euclidean inner product (linear kernel) between these score vectors. This approach allows the linear kernel to quantify the overall similarity between proteins, despite the non-PD nature of the raw BLAST scores.

Proteins with more similar score profiles tend to yield larger inner products under the linear kernel [1], a pattern consistent with observations from BLAST-based comparisons, where higher alignment scores typically reflect stronger biological similarity. The linear kernel thus captures the similarity between two proteins: higher values in the (*i, j*)−th (consequently, (*j, i*)−th) entry in *Q*^(1)^ indicate stronger similarity between proteins *i* and *j*, while lower values suggest weaker similarity.

##### *Q*^(2)^: RBF kernel matrix derived from gene expression data

The gene expression of each protein is represented as a vector, and when the radial basis function (RBF) of each pair of gene expression vectors is computed, the kernel matrix *Q*^(2)^ is constructed. Higher values in *Q*^(2)^ entries indicate greater similarity in gene expression profiles, while lower values suggest weaker similarity [1].

##### *Q*^(3)^: Diffusion kernel matrix derived from protein-protein interaction network

The diffusion kernel is computed from the graph Laplacian matrix associated to the known protein-protein interaction network. Higher values of *Q*^(3)^ entries indicate stronger functional similarity in PPI network, while zero-valued (*i, j*)−th (or (*j, i*)−th) entries means no direct interaction between proteins *i* and *j*.

Thus, for a set of *l* proteins, three *l*×*l* kernel matrices were constructed, corresponding to each data sources shown in Table 1. For *k* ∈ {1, 2, 3}, each *Q*^(*k*)^ is a symmetric and positive semi-definite kernel matrix, where each (*i, j*)−th entry of *Q*^(*k*)^ represents the similarity between proteins *i* and *j* based on the kernel applied to the respective data source. These three kernel matrices are then combined, which we will denote as *Q*^(*c*)^, using the averaging formula [1, 15]:

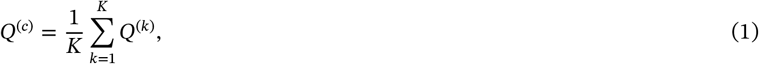

where *K* is the number of kernel matrices. Each row of *Q*^(*c*)^ corresponds to distinct kernel-based feature vector in ℝ^*l*^ for a given protein.

#### 3.2.2. Tracking Persistence of H_0_ and H_1_Topological Features

Topological features were extracted using persistent homology, which captures higher-order structures such as connected components and loops within the kernel-induced similarity space. The kernel matrix *Q* was first transformed into a distance matrix *D*, from which a filtered simplicial complex was constructed. Each protein *i* was treated as a vertex, and edges were added between proteins *i* and *j* if *D*_*ij*_ did not exceed a filtration threshold *ε*. Persistence diagrams were generated using GUDHI [27]. We focused on tracking *H*_0_ and *H*_1_ features since only *H*_0_ and *H*_1_ features were observed in the persistence diagrams, indicating the absence of higher-dimensional topological structures. We monitor the persistence of homological features associated to each protein, stopping when a feature dies, that is, it merges with another feature during filtration.

##### Connected Components, *H*_0_

Each protein *i* begins as an isolated vertex, born at *b*_*i*_ = 0. The corresponding *H*_0_ feature dies at *d*_*i*_ when it merges with another component, marking the loss of independence.

##### Loops, *H*_1_

A loop for protein *i* is born when the last edge completing a cycle appears at *B*_*i*_. It dies at *D*_*i*_ once filled by sufficient 2-simplices (at least *n* − 2 for a cycle of *n* vertices). Proteins however may participate in multiple cycles in the Vietoris–Rips complex so a protein *i* can contribute to several (*B*_*i*_, *D*_*i*_) pairs.

To quantify persistence, we computed the first persistence landscape (FPL), denoted by *λ*_1_(*t*), and its *L*^1^, *L*^2^, and *L*^∞^ norms. Each protein is thus represented by six topological features: three norms of the FPL from *H*_0_, and three norms of the FPL from *H*_1_.

#### 3.2.3. Spectral Features

Spectral features, obtained through eigen-decomposition of the kernel matrices, highlight latent structural patterns in the data. Persistent spectral analysis is implemented via combinatorial Laplacians defined on simplicial complexes across the filtration. At each scale, the eigenvalues of the Laplacian encode connectivity and higher-order structure. Tracking these spectra across the filtration yields a multi-scale descriptor, referred to as persistent spectral analysis, which complements persistent homology.

This perspective complements kernel and topological approaches by emphasizing global organization and clustering tendencies within each dataset. To compute the spectral features for each protein *i*, we define the filtration threshold ϵ_*i*_ as 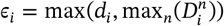 where *d*_*i*_ is the death time of the *H*_0_ associated with protein *i* and 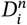 represents the death times of the *H*_1_ features in which protein *i* participates. This choice ensures that the topological signature of each protein fully captures its involvement within the filtration process. In particular, there exist proteins that do not participate in the formation of loops but do contribute to the evolution of connected components. Including *d*_*i*_ ensures that such proteins are not overlooked. On the other hand, for proteins involved in loops, 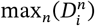 captures the latest topological event in which they are involved. Thus, for each protein *i*, there is a corresponding simplicial complex 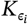 with filtration scale *ϵ*_*i*_ . Let 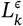 denote the *k*-th combinatorial Laplacian at filtration scale *ϵ*. . We then perform spectral analysis by computing the combinatorial Laplacians 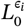 and 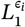 associated with 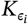. Unlike standard persistent homology, which summarises topological features through birth-death pairs, the spectral perspective captures these structures through the evolution of Laplacian eigenvalues across the filtration, providing a complementary description of connectivity and higher-order organization.

The spectral features of protein *i* are defined as the number of zero eigenvalues of the combinatorial Laplacians 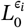 and 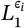 . By [9], dim 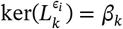 so the multiplicity of the zero eigenvalue of 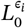 and 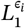 corresponds to the number of connected components and independent 1-dimensional cycles, respectively. The spectral feature of each protein is represented as a vector in ℝ^2^. Figure 2 illustrates the feature extraction component of the pipeline. The completed kernel matrix is first transformed into a distance representation, from which both topological and spectral descriptors (obtained from persistent homology and persistent Laplacian analysis, respectively) are computed and combined into the final feature set.

**Figure 2.**
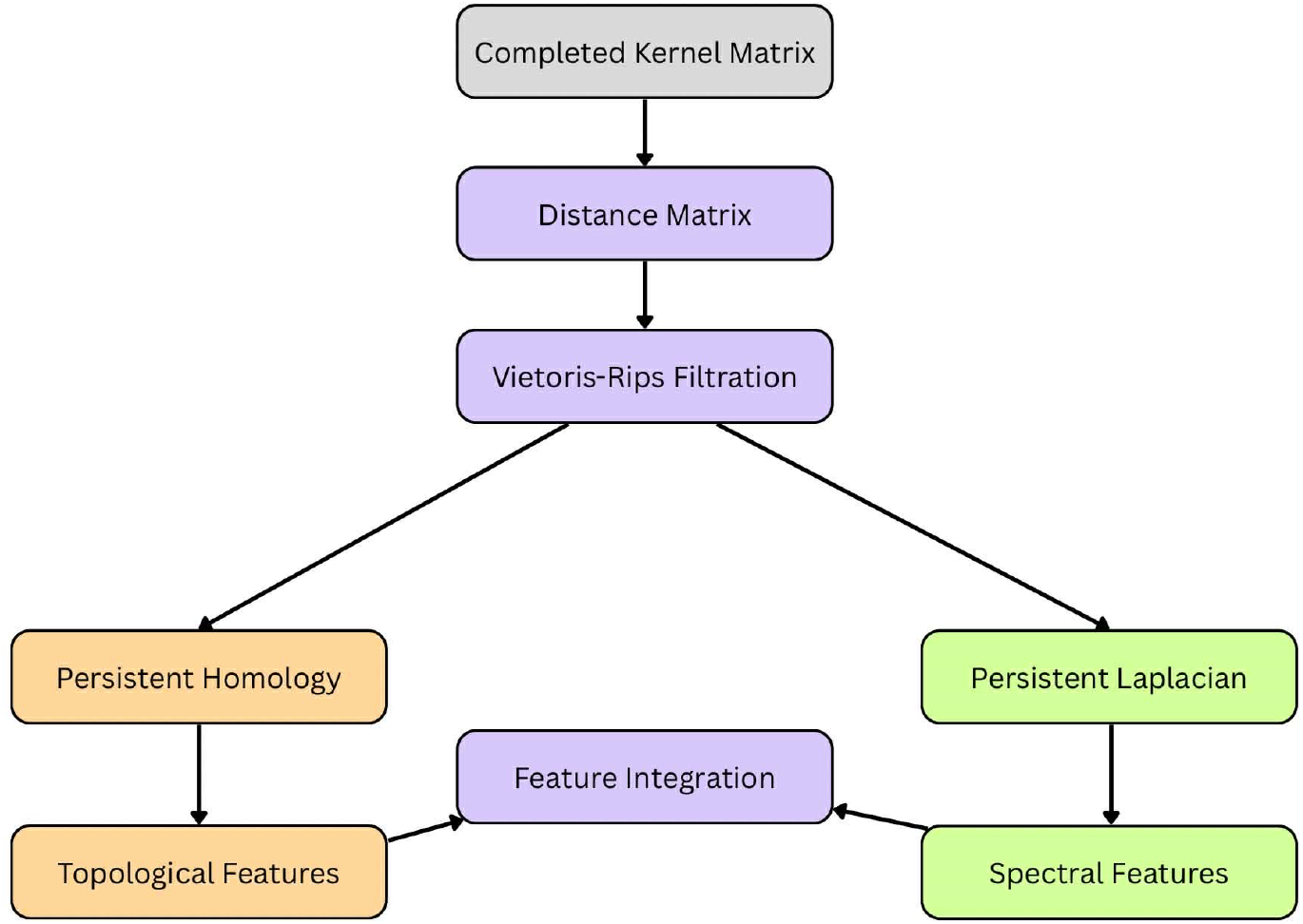
Feature extraction stage of the framework. Persistent homology and persistent Laplacian analysis are applied to the completed kernel matrix to derive topological and spectral descriptors used for classification.

Conceptually, the pipeline can be viewed as a two-stage transformation. The incomplete data are first mapped into a completed similarity representation through kernel matrix completion. This representation is then analyzed at multiple scales via topological and spectral operators, allowing both local similarity structure and global connectivity patterns to be captured in the final feature space. This separation highlights the role of completion as part of representation learning, rather than a standalone preprocessing step.

### 3.3. Classification Task

The computational cost of the pipeline is driven by three main components. Kernel matrices of size *n* × *n* scale as *O*(*n*^2^) in memory, while eigenvalue computations for combinatorial Laplacians and full eigendecomposition scale cubically in the worst case. Persistent homology adds additional cost depending on the size of the simplicial complex. This limits scalability to moderate dataset sizes. Together with the computational limitations of the machine used for the experiments (see Table 2), we were only able to analyze smaller-sized matrices. For a kernel matrix of size 800 × 800, the experiment terminates after seven hours when computing for topological features, and exceeds the computer’s memory when computing for spectral features. To manage this constraint while ensuring a fair evaluation, the 2,318 yeast proteins were randomly divided into four smaller groups—three groups containing 580 proteins each, and one group containing 578 proteins. Each group was formed in such a way that the original proportion of membrane to non-membrane proteins was preserved, allowing the classification task to remain representative of the full dataset.

**Table 2.** Mean computation time for TDA and PSA feature extraction.

| Dimension of $Q$ | Computation Time for Feature Extraction | |
| --- | --- | --- |
|  | Topological Features | Spectral Features |
| $580 \times 580$ | 21 min 30 sec | 33 min |
| $800 \times 800$ | >7 hrs (terminated) | Memory exceeded |
| $2318 \times 2318$ | Memory exceeded | Memory exceeded |

For each group, we performed a classification task to distinguish between membrane and non-membrane proteins using the completed and integrated kernel matrices. To assess the performance of our proposed method alongside the baseline approaches, we used ten-fold cross-validation. In this procedure, each group was divided into ten equal parts: nine parts were used for training the model, and the remaining part was used for testing. This was repeated ten times, with each part serving as the test set once. The classification results were averaged across all folds, and the Receiver Operating Characteristic (ROC) score was used to evaluate classification performance.

## 4. RESULTS AND DISCUSSION

### 4.1. Completion Accuracy

To evaluate how well PCAPSA-MKMC recovered the missing values, we calculated the average correlation distance between the estimated and the original kernel matrices. The metric used is defined as

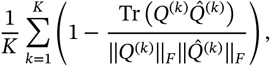

where ‖ ⋅ ‖_*F*_ represents the Frobenius norm, and *Q*^(*k*)^ and 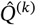 refer to the complete and estimated kernel matrices, respectively. This value was computed for each level of missing data. A lower correlation distance indicates a stronger similarity between the estimated and original matrices, signifying better recovery performance.

In the experiments, the percentage of missing entries was varied from 10% to 90%. As shown in Figure 3, PCAPSA-MKMC consistently achieves lower correlation distances compared to the baseline methods, zero-imputation and PCA-MKMC, even when up to 80% of the data is missing.

**Figure 3.**
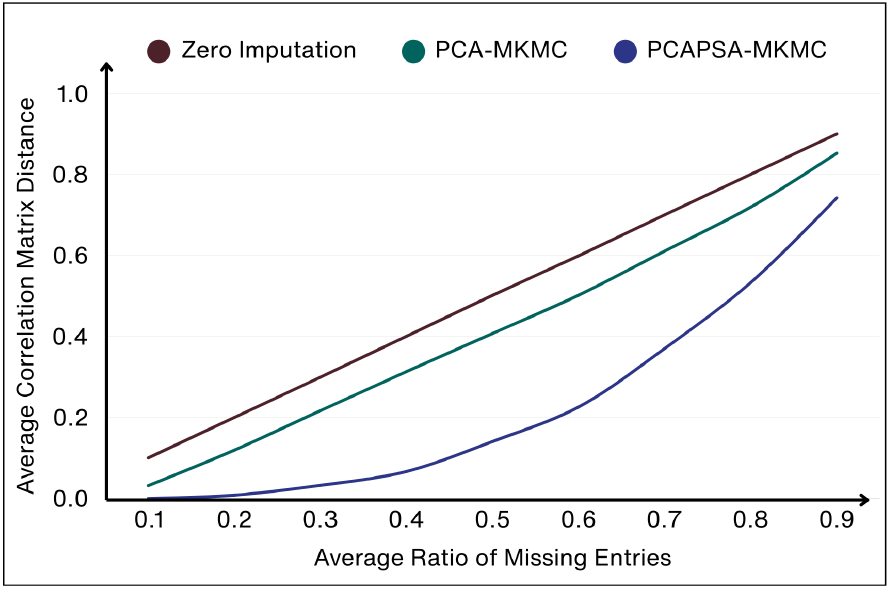
Kernel completion accuracy of zero imputation, PCA-MKMC, and PCAPSA-MKMC across different ratios of missing entries.

Figure 4 presents a series of heatmaps illustrating the impact of varying levels of missing data across three kernel matrices, *Q*^(1)^, *Q*^(2)^, and *Q*^(3)^, on the accuracy of the proposed kernel matrix completion method. Each subplot corresponds to a fixed ratio of missing entries for *Q*^(1)^, ranging from 0.1 to 0.9, while the *x*− and *y*−axes represent the missing ratios for *Q*^(2)^ and *Q*^(3)^, respectively. The color scale encodes the correlation matrix distance, with darker shades indicating lower distances and thus better recovery performance.

**Figure 4.**
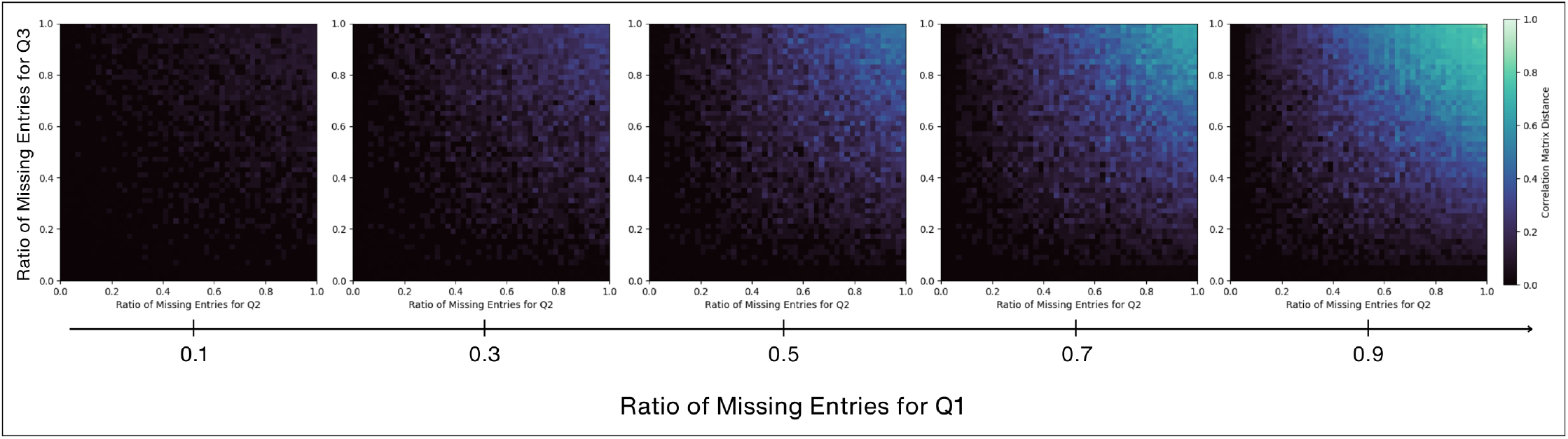
Kernel Completion Accuracy of PCAPSA-MKMC. Each density map represents the average correlation matrix distance, where the ratio of missing entries for *Q*^(1)^ is fixed while the ratio of missing entries for *Q*^(2)^ (horizontal axis) and *Q*^(3)^ (vertical axis).

The results highlight the method’s robustness in handling incomplete data across multiple kernels. When *Q*^(1)^ contains only 10% missing entries, the method consistently achieves low correlation distances across a broad range of missing ratios for *Q*^(2)^ and *Q*^(3)^. This suggests that a nearly complete kernel matrix is sufficient to guide the accurate recovery of highly incomplete ones. As the missing ratio in *Q*^(1)^ increases, a gradual decline in performance is observed, with higher correlation distances appearing when all three kernel matrices exhibit high levels of incompleteness. Notably, the method continues to perform well even when two of the kernel matrices have missing ratios of 80% or more, as long as one of the kernels retains a low missing ratio (e.g., *Q*^(1)^ at 10λ30% missing entries). This suggests that integrating multiple kernel matrices allows the available information in one or two matrices to compensate for missing entries in the others.

### 4.2. Kernel Matrix Completion

To evaluate the performance and robustness of **PCAPSA-MKMC** (Principal Component Analysis with Persistence Spectral Analysis for Mutual Kernel Matrix Completion), we conducted experiments using synthetic missing data scenarios applied to yeast protein datasets shown in Table 1. The three kernel matrices, denoted by *Q*^(1)^, *Q*^(2)^, and *Q*^(3)^, were modified to have different amounts of missing entries, ranging from 10% to 90%, to represent various levels of data sparsity. Missing entries in a kernel matrix were created by choosing certain data and removing their corresponding rows and columns from the matrix. These removed parts were replaced with zeros, but the size of the matrix was kept the same.

To assess the performance of the proposed PCAPSA-MKMC completion method, we compared it against two baseline methods-*zero-imputation* and *PCA-MKMC*. The evaluation was conducted under the following experimental settings:

1. All three kernel matrices were assigned increasing levels of missing data.
2. Two kernel matrices were subjected to varying levels of missing entries (10% to 90%), while the third matrix retained a fixed proportion of missing data across all trials.

### 4.3. TDA

We examined the topological properties of the matrices *Q*^(1)^, *Q*^(2)^, *Q*^(3)^, and *Q*^(*c*)^. Each kernel matrix was constructed from a random subset of yeast proteins. Consequently, for each matrix, we obtained four corresponding submatrices that represent independent random samplings of the dataset. For each submatrix, we computed persistence diagrams that illustrate the birth and death of topological features across a range of filtration values. These diagrams summarize how connected components and loops evolve as the filtration threshold increases. To quantify these observations, we extracted the Betti numbers *β*_0_ and *β*_1_, representing the number of connected components and one-dimensional holes, respectively, at each filtration step. Figures 5 and 6 illustrate the persistence diagrams and Betti number counts of each subgroup of the kernel matrix *Q*^(*c*)^, respectively.

**Figure 5.**
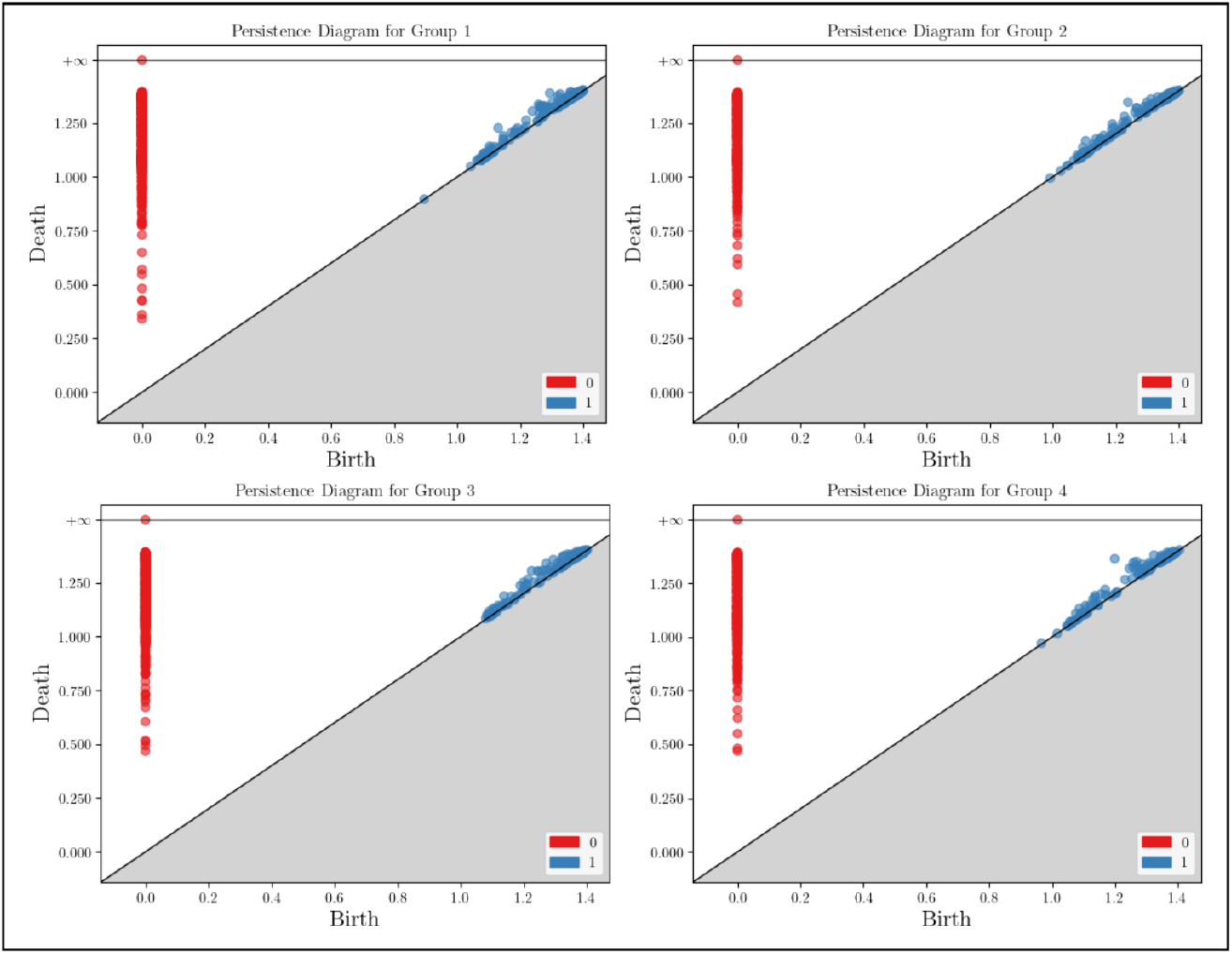
Persistence diagrams corresponding to each subgroup within the kernel matrix *Q*^(*c*)^, showing the birth and death of topological features across filtration.

**Figure 6.**
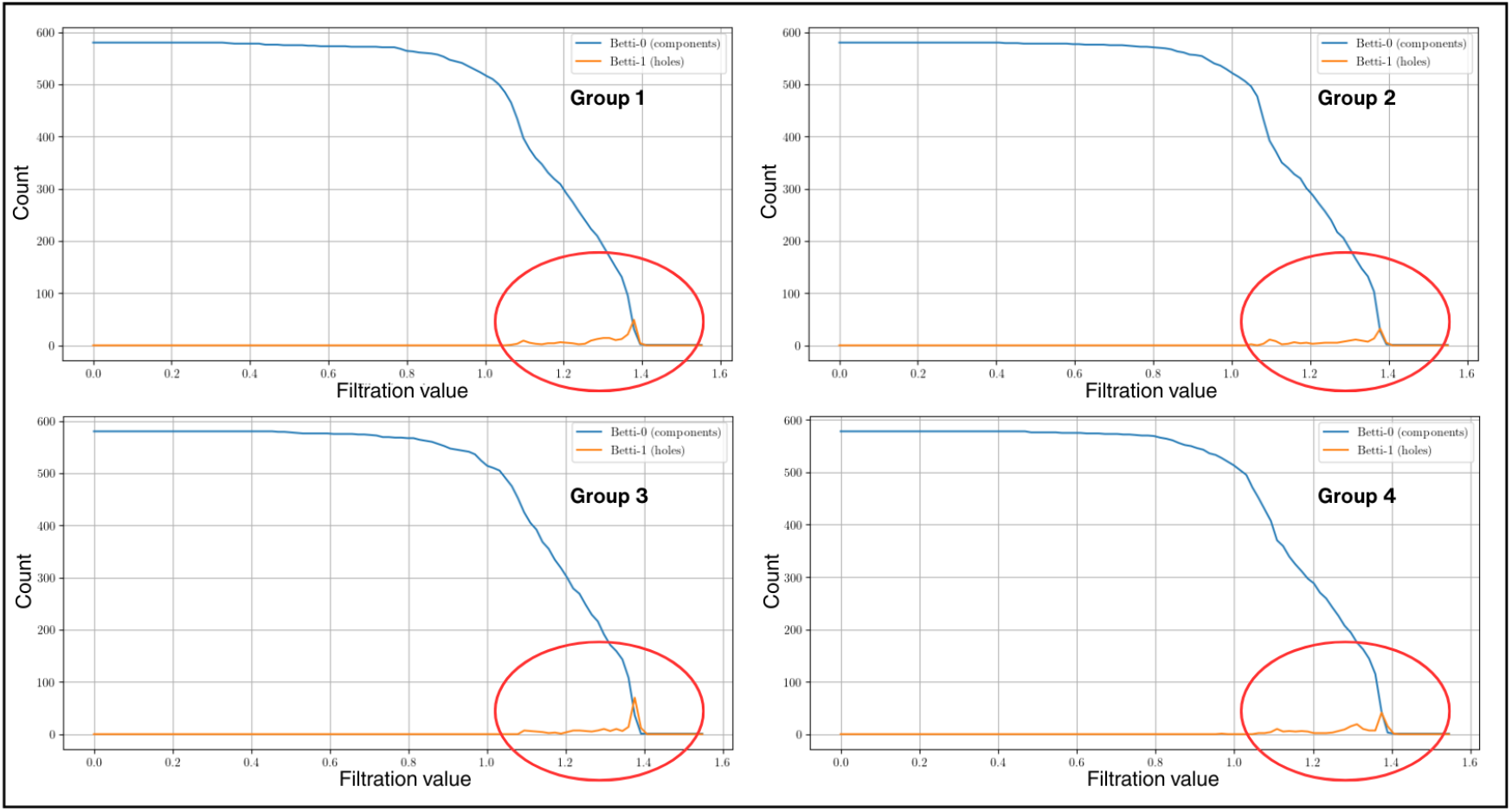
Betti number counts for each subgroup within the kernel matrix *Q*^(*c*)^, showing the number of connected components (*β*_0_) and one-dimensional holes (*β*_1_) across varying filtration values.

Despite the randomness in sampling, the Betti number distributions—computed from the persistence diagrams—remained consistent across the subgroups. In particular, the number of connected components (*β*_0_) and one-dimensional holes (*β*_1_) showed similar patterns across filtration. As the filtration value increased, *β*_0_ decreased as components merged, while *β*_1_ features appeared temporarily, indicating the presence of loops. These trends suggest that the overall data connectivity and geometric structure are preserved across the different groups, and that the topological features captured by each kernel matrix are stable under random partitioning. This consistency across subgroups suggests that the kernel-induced geometry preserves meaningful structural information regardless of the specific composition of the subset. As such, the use of TDA in conjunction with kernel matrices forms a robust framework for analyzing the topological characteristics of the protein dataset, even under constrained computational settings that necessitate partitioning.

Interestingly, we observed a noticeable spike in the *β*_1_ counts for the RBF kernel around a filtration value of approximately 1.25 shown in Figure 7, in contrast to the more stable trends exhibited by the linear, diffusion, and mixed kernels. This spike suggests that the RBF kernel may be capturing a large number of loosely connected protein clusters that momentarily form one-dimensional loops. Given that these features appear and vanish rapidly, they are likely indicative of topological noise rather than meaningful structure.

**Figure 7.**
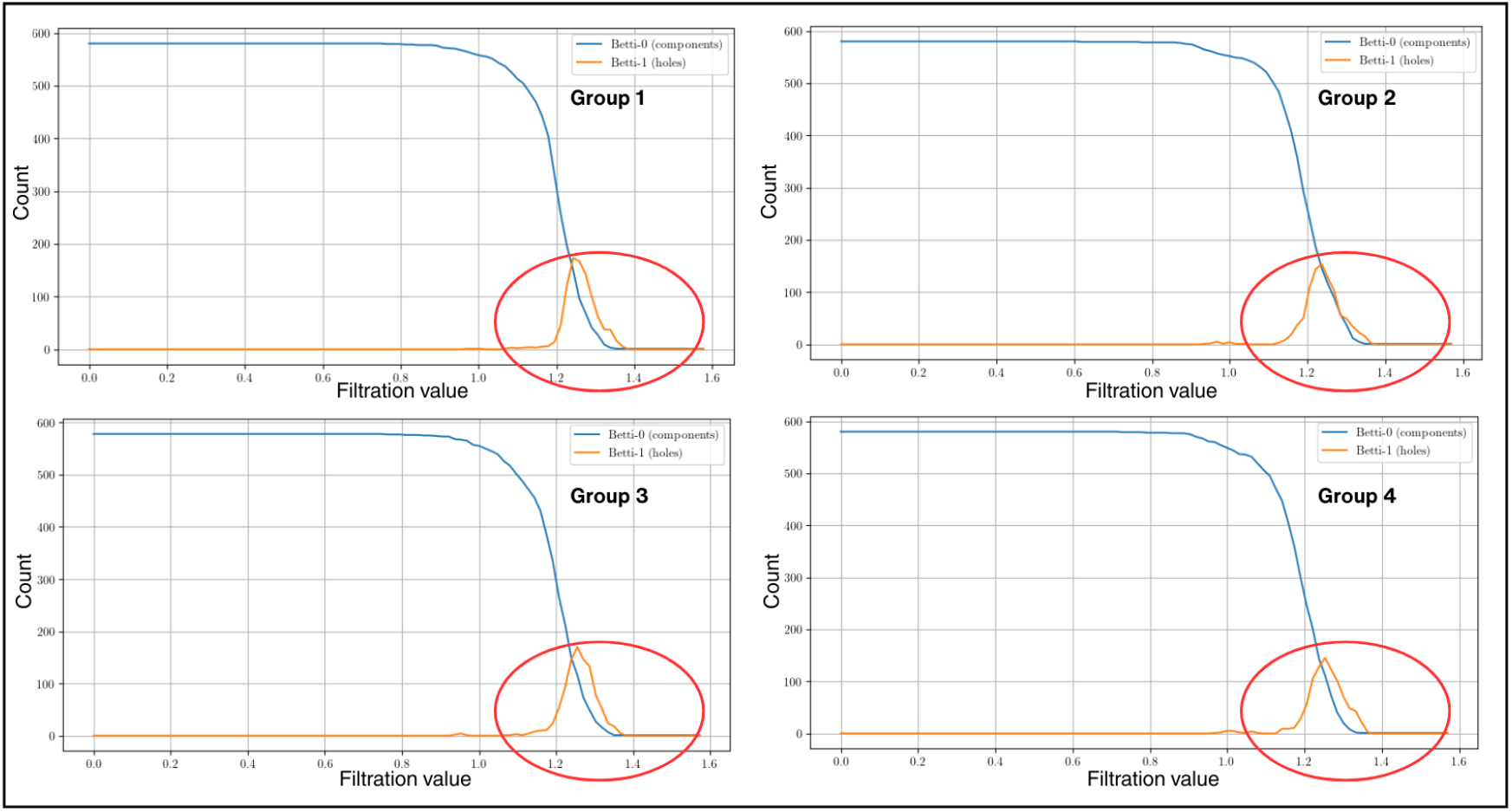
Betti number counts for each subgroup within the kernel matrix *Q*^(2)^ (RBF kernel), showing the number of connected components (*β*_0_) and one-dimensional holes (*β*_1_) across varying filtration values.

### 4.4. Classification Performance

We assessed the performance of the SVM model using individual kernel matrices, their completed form, and their integration with TDA and PSA. Classification accuracy was evaluated using the area under the ROC curve. Table 3 summarizes the ROC scores across protein groups, along with their corresponding standard deviations (SD). Among the three individual kernels, the linear kernel matrix *Q*^(1)^ achieved the highest mean ROC score of 0.71. Using the completed kernel matrix *Q*^(*c*)^ alone gives an average ROC score of 0.8046. This suggests that the integration of multiple kernel matrices captures relevant structural relationships between protein features. While the completed kernel matrix demonstrates good results on its own, further improvements in classification performance were observed when incorporating the additional contributions of PSA and TDA. When PSA and TDA are used together, we observed improvements in classification classification performance, resulting in higher ROC scores. Notably, combining the completed kernel matrix with TDA produced the highest ROC score, while integrating PSA alone also demonstrated good performance. Overall, the combined kernel *Q*^(*c*)^ proved most effective and robust. Adding TDA consistently improved performance, especially for *Q*^(*c*)^, while PSA yielded moderate gains. However, combining both TDA and PSA with *Q*^(*c*)^ did not further improve results, suggesting redundancy between the two. Notably, *Q*^(2)^+TDA showed the highest variability, likely due to topological noise from short-lived loops in the RBF kernel.

**Table 3.** Classification performance (ROC scores) across protein groups of individual and combined kernels with TDA and PSA, for 20% missed kernel data. The boldfaced values correspond to the largest mean ROC score per feature.

| Feature | Group 1 | Group 2 | Group 3 | Group 4 | Mean | SD |
| --- | --- | --- | --- | --- | --- | --- |
| $Q^{(1)}$ | 0.6600 | 0.7307 | 0.6632 | 0.7870 | 0.7102 | 0.0607 |
| $Q^{(1)}+TDA$ | 0.6939 | 0.7966 | 0.7144 | 0.7295 | 0.7336 | 0.0385 |
| $Q^{(1)}+PSA$ | 0.7364 | 0.7942 | 0.7568 | 0.7849 | **0.7681** | 0.0229 |
| $Q^{(1)}+TDA+PSA$ | 0.6901 | 0.7392 | 0.7165 | 0.6987 | 0.7111 | 0.0188 |
| $Q^{(2)}$ | 0.5466 | 0.5891 | 0.6097 | 0.6480 | 0.5984 | 0.0423 |
| $Q^{(2)}+TDA$ | 0.7146 | 0.5195 | 0.5117 | 0.5647 | 0.5776 | 0.0816 |
| $Q^{(2)}+PSA$ | 0.6641 | 0.6859 | 0.6987 | 0.7092 | **0.6895** | 0.0168 |
| $Q^{(2)}+TDA+PSA$ | 0.7118 | 0.6967 | 0.6765 | 0.6359 | 0.6802 | 0.0285 |
| $Q^{(3)}$ | 0.6475 | 0.6564 | 0.7737 | 0.6945 | 0.6770 | 0.0298 |
| $Q^{(3)}+TDA$ | 0.7960 | 0.7358 | 0.7436 | 0.7685 | **0.7610** | 0.0236 |
| $Q^{(3)}+PSA$ | 0.7508 | 0.7513 | 0.7440 | 0.7927 | 0.7597 | 0.0193 |
| $Q^{(3)}+TDA+PSA$ | 0.7045 | 0.6838 | 0.7045 | 0.7362 | 0.7073 | 0.0187 |
| $Q^{(c)}$ | 0.7934 | 0.8122 | 0.7961 | 0.8167 | 0.8046 | 0.0116 |
| $Q^{(c)}+TDA$ | 0.8409 | 0.8529 | 0.8774 | 0.8814 | **0.8632** | 0.0194 |
| $Q^{(c)}+PSA$ | 0.8050 | 0.8344 | 0.8388 | 0.8400 | 0.8296 | 0.0165 |
| $Q^{(c)}+TDA+PSA$ | 0.8099 | 0.8315 | 0.8362 | 0.8391 | 0.8298 | 0.0138 |

Although our highest ROC does not exceed the best value reported in [1], we note that their method assumes complete data and focuses on learning optimal kernel weights in a supervised framework. In contrast, our work considers the incomplete-data scenario and incorporates kernel matrix completion together with topological and persistent spectral features. From this perspective, our proposed framework remains competitive while extending the analysis to incomplete heterogeneous data.

While the proposed framework does not aim to outperform methods designed for fully observed data, the results indicate that it achieves competitive performance under incomplete data conditions. In particular, the use of topological descriptors together with completed kernel matrices yields consistent improvements over baseline kernel approaches, highlighting the importance of incorporating structural information when learning from partially observed similarities.

## 5. CONCLUSION

In this work, we introduced PCAPSA-MKMC, a novel framework for kernel matrix completion and pattern analysis that addresses the challenge of learning from incomplete, high-dimensional biological data. By incorporating topological and spectral descriptors into the matrix recovery process, we show that PCAPSA-MKMC outperforms traditional methods, achieving lower correlation distances between the estimated and original kernel matrices, even under extreme sparsity conditions of up to 80% missing data.

The primary strength of this approach lies in its unified representation learning through cross-modal complementarity. While the completed kernel matrix serves as a robust foundation, the incorporation of TDA and PSA captures complementary structural patterns that were often overlooked by traditional methods. In the membrane versus non-membrane classification task, the inclusion of TDA was particularly effective, yielding a peak ROC score of 0.8632, highlighting the role of topological features in refining complex decision boundaries.

The proposed framework highlights the role of combining similarity-based, topological, and spectral representations when learning from incomplete structured data. From a pattern recognition and computational biology perspective, this work provides a more structured approach to manifold learning and feature extraction in noisy environments, and a systematic pipeline for handling incomplete kernel matrices without resorting to naive imputation methods, such as zero- and mean-imputation. This is especially relevant for practitioners working with multi-modal data sources where reconciliation is required across varying levels of data quality. However, a notable limitation is feature redundancy, as our findings indicate that PSA and TDA do not always yield additive gains when used together. This suggests that these descriptors may capture overlapping structural information, highlighting a need for more sophisticated fusion strategies to maximize the utility of each modality. In addition, the current framework does not include a direct comparison with recent graph-based or deep learning approaches, which may offer complementary advantages in large-scale settings.

While our current results are promising, several directions for development remain, particularly regarding scalability and computational efficiency. When the matrices exceeded 580 × 580 dimensions, the memory demands outpaced available resources. This computational cost arises from the use of persistent homology. The next steps would be to refine the framework using sparse TDA and adaptive fusion, which will allow the system to weigh different data types more effectively. Aside from producing a more efficient algorithm, we also aim to make TDA more accessible and practical for the high-volume data demands of biological and pattern recognition fields. Extending the framework to larger datasets and to other pattern recognition tasks involving incomplete and heterogeneous data will be important in showing how it can be applied more broadly.

From a computational perspective, the main bottlenecks arise from the construction of simplicial complexes and the associated spectral computations, which scale with both the number of data points and the density of the complex. While this limits the current implementation to moderate-sized datasets, it also suggests clear directions for improvement, including sparse filtrations, approximate spectral methods, and sub-sampling strategies that preserve topological structure.

In summary, the proposed framework addresses the challenge of learning from incomplete heterogeneous data by integrating similarity-based, topological, and spectral representations within a unified pipeline. By combining kernel matrix completion with multi-scale feature extraction, the approach preserves structural information and improves classification performance under incomplete data conditions, demonstrating the effectiveness of structured representations for multi-modal data. This shows that the framework can be used for pattern recognition tasks involving incomplete and multi-modal data.

## DATA AVAILABILITY

The protein dataset used in this study is publicly available at https://noble.gs.washington.edu/proj/sdp-svm/.

## ACKNOWLEDGMENTS

Numerical results of this work were generated at the Computational Research Laboratory of the Institute of Mathematics, University of the Philippines Diliman. MVV acknowledges support from the DSI-NRF Centre of Excellence in Mathematical and Statistical Sciences (CoE-MaSS). Opinions expressed, and conclusions arrived at are those of the authors and are not necessarily to be attributed to the CoE-MaSS.

## Notes

https://noble.gs.washington.edu/proj/sdp-svm/?fbclid=IwY2xjawQtcHBleHRuA2FlbQIxMABicmlkETFsSGlyVHcydElzRzVndjlwc3J0YwZhcHBfaWQQMjIyMDM5MTc4ODIwMDg5MgABHmbEBfdpZNDXh0XSSHhFdZmI7plpD80EkhGKTi7nv3Y2vLEKKTqbeKY4aInT_aem_UvjgGgP7HaH3ObKLQ4O8aQ

### Competing Interest Statement

The authors have declared no competing interest.

